# Chromosomal inversions can limit adaptation to new environments

**DOI:** 10.1101/2022.05.02.490344

**Authors:** Marius Roesti, Kimberly J. Gilbert, Kieran Samuk

## Abstract

Chromosomal inversions are often thought to facilitate local adaptation and population divergence because they can link multiple adaptive alleles into non-recombining genomic blocks. Selection should thus be more efficient in driving inversion-linked adaptive alleles to high frequency in a population, particularly in the face of maladaptive gene flow. But, what if ecological conditions and hence selection on inversion-linked alleles change? Reduced recombination within inversions could then constrain the formation of optimal combinations of pre-existing alleles under these new ecological conditions. Here, we outline this idea of inversions limiting adaptation and divergence when ecological conditions change across time or space. We reason that the benefit of inversions for local adaptation and divergence under one set of ecological conditions can come with a concomitant constraint for adaptation to novel sets of ecological conditions. This limitation of inversions to adaptation may also provide one possible explanation for why inversions are often maintained as polymorphisms within species.

## Background

In evolutionary biology, there is a common notion that chromosomal inversions facilitate adaptation and divergence. Inversions create different physical arrangements of a genomic region, which often leads to non-viable gametes when recombination between these arrangements occurs (Sturtevant et al. 1936; Navarro et al. 1997). As a result, realized recombination between different inversion arrangement types is strongly reduced at the population level, and alleles within one arrangement type become strongly linked and can behave similarly to a single allele of large selective effect. Selection should thus be more efficient in maintaining sets of inversion-linked alleles if they are adaptive and driving them to high frequency in a population, particularly in the face of maladaptive gene flow (Rieseberg 2001). Indeed, theory suggests that local adaptation of a population can be achieved more readily when multiple, locally adaptive alleles are contained within the same inversion arrangement type (Kirkpatrick & Barton 2006; Feder & Nosil 2009; Charlesworth & Barton 2018).

Consistent with the idea of inversions facilitating local adaptation and divergence, one inversion arrangement type is often found at a relatively high frequency within populations, and populations from different habitats often differ strongly in their frequency of arrangement types (e.g., Wellenreuther & Bernatchez 2018; Faria et al. 2019). However, recent work has highlighted that reduced recombination between inversion arrangement types can hinder the purging of unconditionally (i.e., environment-independent) deleterious mutations, such as premature stop codons or recessive lethals (Berdan et al. 2021; Jay et al. 2021). The accumulation of such deleterious mutations may thus counteract the adaptive potential of inversions for local adaptation. For recessive deleterious variants, the reduction in recombination resulting from inversions may also lead to patterns of associative overdominance, where there is an apparent heterozygous advantage due to masked deleterious variants (Gilbert et al. 2020). This type of balancing selection or the combination of both beneficial and unconditionally deleterious variants within a single inversion provide possible explanations for why inversions may often be maintained as polymorphisms within species (Berdan et al. 2021; Jay et al. 2021).

Another limitation to adaptation from inversions could occur when selection favors new combinations of existing inversion-linked alleles. This can happen due to temporally or spatially varying selection. When selection changes in direction, pre-existing inversion arrangements could pose a constraint to further adaptation because recombination cannot build optimal combinations from pre-existing alleles bound within inversions. The idea that inversions could constrain selection from favoring optimal allele combinations at inversion-linked adaptive loci is distinct from the accumulation of unconditionally deleterious mutations and could represent an important explanation for the evolution and maintenance of chromosomal inversions among natural populations.

## The adaptive limitation hypothesis of inversions

Mounting empirical evidence suggests that standing genetic variation is the main source of genetic variation for the early phases of adaptation in nature (e.g., Renaut et al. 2011; Jones et al. 2012; Lescak et al. 2015; Haenel et al. 2019; Lai et al. 2019; Chaturvedi et al. 2021; Louis et al. 2021; Owens et al. 2021; Whiting et al. 2021; see also Barrett & Schluter 2008; Messer & Petrov 2012; De Lafontaine et al. 2018). Whether and how rapidly a population can adapt to a new ecological challenge therefore depends on how efficiently selection can reshape pre-existing alleles into new optimal combinations. Inversions may limit such genetic reshaping.

Imagine a scenario where each of two different inversion arrangements contains alleles that are beneficial in one habitat type and maladaptive in another habitat type. Then, a new third habitat type becomes available favoring a novel combination of these alleles from the two arrangements. The lack of recombination between the arrangement types will hinder reshaping of optimal allele combinations and hence can limit rapid adaptation into the new habitat **(Figure 1A)**. Similarly, if ecological conditions and thus selection changes for one or both of the initial populations, the lack of recombination of pre-existing alleles between arrangement types could impede adaptation compared to when adaptive alleles are not inversion-linked and thus free to recombine. Both of these scenarios, a novel habitat appearing or an existing habitat changing, are representative of multitudes of real-world scenarios, which can drastically alter the direction of natural selection.

**Figure 1.**
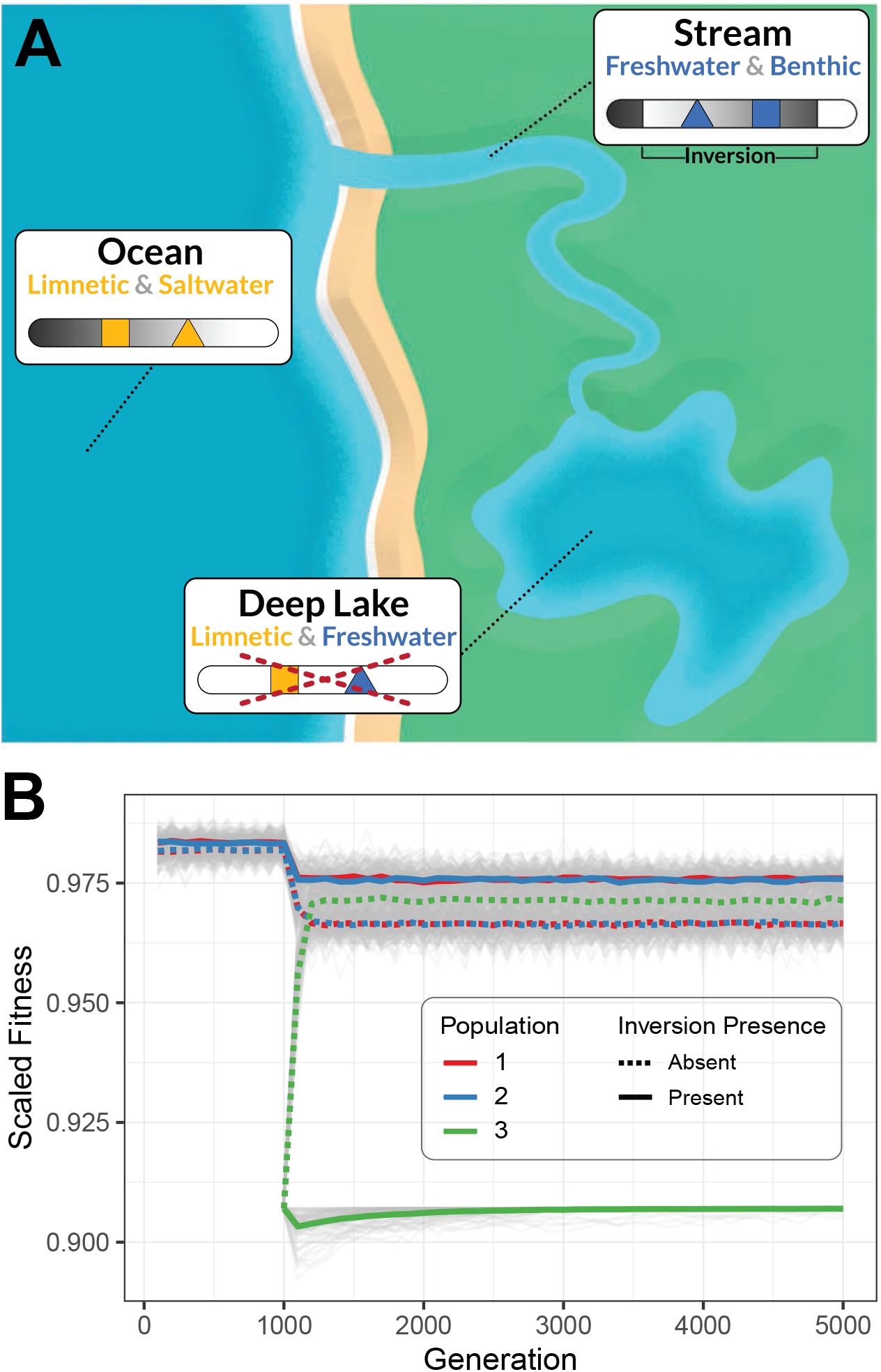
Exemplary scenario and simulation of how inversions can limit adaptation to new environments. **(A)** In this exemplary scenario, inversion-linked alleles at two biallelic loci confer adaptation to two different original habitats in an aquatic organism: saltwater and limnetic alleles (ocean habitat) vs. freshwater and benthic alleles (stream habitat). Such an inversion will limit optimal adaptation into a novel third habitat (deep lake) that requires the combination of freshwater and limnetic alleles. **(B)** Results from forward-in-time simulations using SLiM (Haller & Messer 2019), based on the scenario outlined in (A). Population 3 forms at generation 1000 and represents the novel deep lake habitat, which in the absence of an inversion can be successfully colonized, reaching relatively high population fitness in the face of migration-selection balance (dashed lines). In the presence of the inversion (solid lines), however, fitness is reduced in the novel habitat as optimal adaptation is prevented from the alleles locked within the inversion. In these simulations, each allele has an equal selective and thus fitness effect, being beneficial in one of the two original habitats and detrimental in the other, with *s* = +/-0.1. In population 3, the allele favored (+s) in population 1 at the first locus and the allele favored in population 2 at the second locus are favored. Adaptation of each population is expressed as the mean population relative fitness scaled against the maximum possible relative fitness based on the known optimal genotypes (i.e., a scaled fitness of 1 represents optimal adaptation of a population). Thick lines in color indicate the mean scaled fitness of 100 replicate simulations (gray lines). De novo mutation or double-crossovers were not considered in these simulations. See the Supplementary Materials for further details on the simulations as well as alternative scenarios and parameter combinations tested, including a polygenic model.

To illustrate this idea, we explored whether inversions limit adaptation in forward-time individual-based simulations mimicking these two scenarios. Simulations begin with a two-deme model in which each of two populations adapts to a distinct environment. Individuals are diploid and possess a genome with two loci, each with two alleles conferring adaptation to either one of the two environments, respectively (i.e., these loci are under divergent selection between the populations). Populations exchange migrants and thus alleles throughout the duration of the simulation. In one scenario, we then introduce a new third habitat which can be colonized **(Figure 1B, Fig. S1)**. Alternatively, in a second scenario, we change the environment for one of the existing populations **(Fig. S2)**. In both cases, novel selective pressure now favors a new combination of alleles at the two loci: selection favors the allele adaptive in population 1 at one locus, and the allele adaptive in population 2 at the other locus. We ran these simulations both with and without an inversion that captured one of the two sets of alleles adaptive in either one of the two initial populations as an arrangement. Overall, these simulations confirm our intuition that an inversion can limit adaptation to a new adaptive optimum compared to simulations without inversions where optimal combinations of pre-existing alleles can be created easily via recombination **(Figure 1B, Figs. S1 and S2**).

These simulations are intentionally simplified and do not explore the full range of conditions under which an inversion can limit adaptation to changing adaptive optima. Yet these results do demonstrate that, in principle, inversions can limit rapid local adaptation and hence adaptive divergence between populations. Although we placed reciprocally adaptive/maladaptive alleles within alternative inversion arrangements, a similar (albeit weaker) effect could be generated by an inversion that was polymorphic but unrelated to the change in selection (e.g., because it contains a recessive lethal allele). In this case, a reduction in average recombination in the inverted region would result in the limitation of adaptation via standard Hill-Robertson interference (Hill & Robertson 1966).

Our described constraint of reduced recombination at inversions for adaptation is conceptually related to the long-standing idea for why asexual reproduction is particularly disadvantageous when environments change frequently over time or space. That is, maladaptive genetic associations built by past selection or brought to a different environment through migration cannot be rebuilt into favorable combinations in the absence of recombination as it is the case in asexually reproducing organisms (Maynard Smith 1978; Otto 2009). Another conceptual parallel can be drawn to the constraint described previously for pleiotropy, where a single gene affects multiple traits and may therefore hinder the evolution of optimal trait combinations under varying ecological conditions (Cheverud 1984; Pavlicev & Cheverud 2015). These conceptual parallels between asexual reproduction, pleiotropy, and inversions can help explain how the absence of recombination can constrain adaptive evolution, yet the dynamics of inversions are unique and worth special consideration since recombination is only reduced in individuals carrying both arrangement types (heterozygotes).

### Outcomes and future investigations

There are several ways by which the adaptive limitation of inversions could resolve itself genetically. Gene conversion or double-crossovers could allow for rare genetic exchange (gene flux) between inversion arrangement types, thereby allowing for the build-up of the combinations of pre-existing alleles that are favorable under changed ecological conditions. *De novo* mutations in pre-existing inversion arrangements as well as in other regions of the genome could also build newly favored allele combinations. While both of these routes could resolve the limitation that inversions can pose to adaptation, they will necessitate longer wait times than a normally recombining genomic region. Moreover, these considerations emphasize the need for a greater appreciation of the genetic variation within – and not only between – inversion arrangement types.

The here-described idea of how inversions may limit rapid adaptation to changing ecological conditions seems compatible with observations in nature. For instance, QTL underlying trait variation that is important for adaptive divergence across a major habitat transition have been mapped to chromosomal inversions in populations of threespine stickleback fish and littorina snails (stickleback: Peichel & Marques 2017; Liu et al. 2021; littorina: Koch et al. 2021). However, both of these species have recently been exposed to new niches imposing novel selection pressures, possibly favoring novel combinations of these inversion-linked QTL (stickleback: e.g., Bell & Foster 1994; Reid et al. 2021; littorina: Morales et al. 2019).

Direct tests of how frequently inversions pose a limit to adaptation in nature will be challenging, especially because genetic variants within inversions are in strong linkage and therefore difficult to assay individually. A promising yet challenging approach would be to unlock inversion-linked genetic variants by flipping one arrangement using Crispr/Cas9-induced double strand breaks, thereby restoring collinearity and thus recombination between different inversion arrangement types (Schmidt et al. 2020). This would subsequently allow for estimating how selection targets individual alleles that were previously inversion-linked. An adaptive constraint of inversions would be implicated if selection targeted some of the previously linked alleles within an arrangement type in the opposite direction within the given ecological context. Another less direct test of the adaptive limitation hypothesis of inversions could use QTL mapping of ecologically-important trait variation (analogous to a QTL sign test; Orr 1998). An adaptive constraint of an inversion may be implicated if the trait effects of some within-inversion QTL were reversed to what would be expected under optimal adaptation. Finally, if inversions are indeed hotspots of adaptive loci, one might expect that the genetic variation unique to the distinct arrangements of a (single) large inversion is unlikely to play a key role in the rapid diversification of a taxon into many niches, and may even pose a constraint for such adaptive radiations. This constraint could be counteracted by the existence of several inversions if each inversion captures a combination of alleles that allows successful adaptation in the face of gene flow across independent environmental axes.

## Conclusion

While an inversion can link unique adaptive allele combinations into non-recombining genomic blocks (haplotypes) and thereby favor local adaptation under one set of ecological conditions, this benefit may come with a concomitant constraint in adaptation to a novel set of ecological circumstances. Indeed, inversions linking unique allele combinations into distinct haplotypes may also be prone to be maintained as polymorphisms within species under spatially and/or temporally varying selection. While searching for evidence of such adaptive limitations imposed by inversions in nature will be challenging, further investigation of this phenomenon will broaden our understanding of the processes shaping diversity across variable environments and during rapid adaptive radiations.

## Acknowledgements

We would like to thank Gregory Owens, Stephan Peischl, Sam Yeaman, and members of the Peichel lab for stimulating and helpful discussions. MR was funded by the University of Bern. KJG was funded by a Swiss National Science Foundation Ambizione grant (PZ00P3_185952).

## Supplementary Materials

### Supplementary Methods

#### Description of simulations

Our illustrative simulations were performed using SLiM version 3.7.1 (Haller and Messer 2019). The code to produce all the simulation results as well as to create the plots shown in the manuscript is available at https://github.com/ksamuk/inversion_constraint. A brief description of the simulation follows.

#### General simulation structure

Each simulation had the following basic structure. Two populations, population 1 and population 2 (hereafter p1 and p2), are initialized in the first generation. Each population is composed of 2500 hermaphroditic individuals, with each individual having a diploid “genome” composed of three loci: two fitness-related loci, and one “inversion” locus. Each of the two fitness loci have two alleles: the “m1” allele and the “m2” allele, each favored in p1 and p2 respectively. Each allele has a selection coefficient of *s* = 0.05 in its home population, and a coefficient of -*s* (i.e., −0.05) in the alternate population (i.e., symmetrical divergent selection). All alleles had dominance coefficients of *h* = 0.5 (i.e., pure additivity). The inversion locus similarly had two alleles, “m3” and “m4”, corresponding to the genomic rearrangements in p1 or p2 respectively. No other mutations were possible in the simulation (i.e., all adaptation occurred from standing variation).

Because of the simplified genomic architecture, we set baseline recombination rates at a value of 1e-2 (SLiM units) in order to observe sufficient recombination over the course of the simulation. We modelled recombination suppression by the inversion after the example in SLiM manual, i.e., if an individual is heterozygous at the inversion locus, recombination is suppressed at all three loci (the fitness loci and the inversion locus). Otherwise, recombination proceeds at the baseline rate.

The two populations exchanged migrants at a rate of *m* = 0.01 (symmetrical gene flow). The simulations were run for 5000 generations. At each generation, we output the mean relative fitness of each population, scaled against the maximum possible fitness based on the known optimal genotypes. To examine the effect of the inversion, we ran simulations with and without the inversion active. We ran 100 replicates of each simulation, and then processed and plotted the output from SLiM using R version 4.1.2 (R Core Team, 2021) and the tidyverse package (Wickham et al. 2019).

Using this core structure, we simulated three different scenarios:

##### (i) “Novel environment” scenario

In this scenario, the simulation proceeds as before, but at *t* = 1000 generations, a third population, population 3 (hereafter p3), is founded with half its individuals sourced from p1 and the other half from p2. In p3, at the first fitness locus, m1 allele has a selection coefficient of *s* = 0.05 and m2 has a selection coefficient of *s* = −0.05. At the second fitness locus, the m2 allele has a selection coefficient of *s* = 0.05 and m1 has a selection coefficient of *s* = −0.05. As such, the optimal genotype in p3 is m1/m1 at the first locus, and m2/m2 at the second locus, i.e., an intermediate between p1 and p2. After the initial founding event, gene flow between all populations continued at the rate of *m* = 0.01.

##### (ii) “Environmental change” scenario

This scenario is similar to the novel environment scenario above, but instead of a new population being founded at *t* = 1000, only two populations still exist and instead the selection coefficients in p2 shift to those described for p3 above (i.e., the optimal genotype is intermediate between the original p1 and p2). All other parameters, including migration rate, remain unchanged.

##### (iii) Polygenic scenario

To explore the effect of a more complex genetic architecture, we simulated the novel environment scenario with a genome containing 101 loci: 100 fitness-related loci and one inversion locus, or 40 fitness-related loci, 60 neutral loci, and one inversion locus. The simulations were otherwise identical.

#### Basic exploration of parameter space

While not our primary goal, we explored the robustness of our results by varying the strength of selection (*s*) and migration rate (*m*) at 0.01, 0.05, and 0.1 in the novel environment scenario. All simulations had qualitative similar results, i.e., the inversion acts as a constraint on adaptation.

## Supplemental Figures

**Figure S1.**
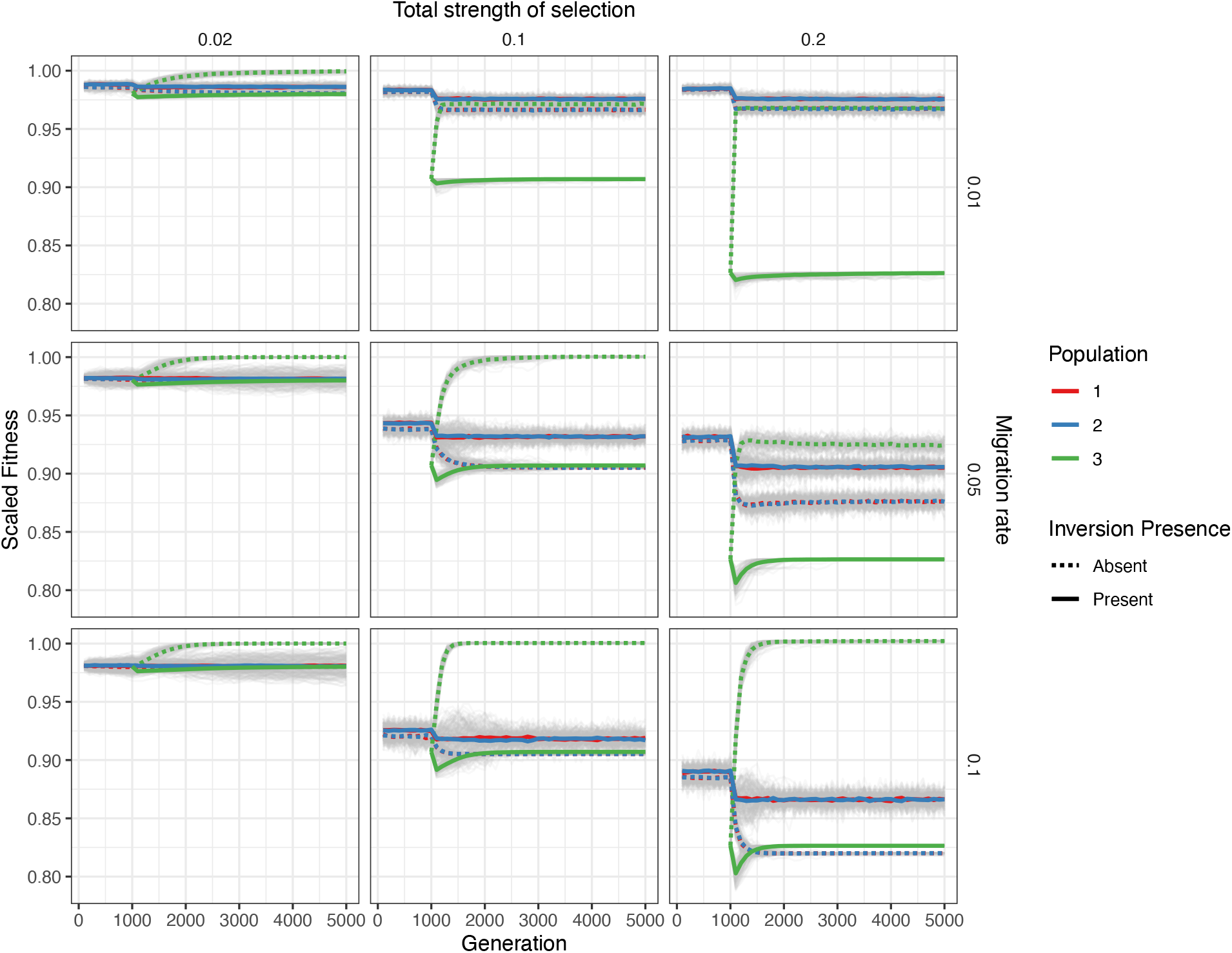
Simulation results for the three populations in the “novel environment” scenario across nine parameter combinations. Each panel depicts the change in scaled mean fitness of a simulated population (colors) over time in the presence or absence of a chromosomal inversion (solid and dashed lines, respectively) under a single parameter combination. Each column depicts simulations performed at different total strengths of selection (the sum of the magnitude of all selection coefficients across all loci in any of the given environments). Each row depicts simulations performed at different migration rates. Note that under each parameter combination, the presence of an inversion limits adaptation of population 3 into the novel habitat (i.e., the dashed green line is always above the solid green line). It is also interesting to note that in both population 1 and population 2, fitness is reduced less in the presence of the inversion as compared to the absence of the inversion. This is because when the inversion is present, population 3 cannot reach its optimal genotype, so the genotypes adaptive in both population 1 and population 2 are at a higher prevalence and thus cause less maladaptive gene flow back into population 1 and population 2. For further details on the simulations, see Supplementary Materials and the caption of Fig.1B.

**Figure S2.**
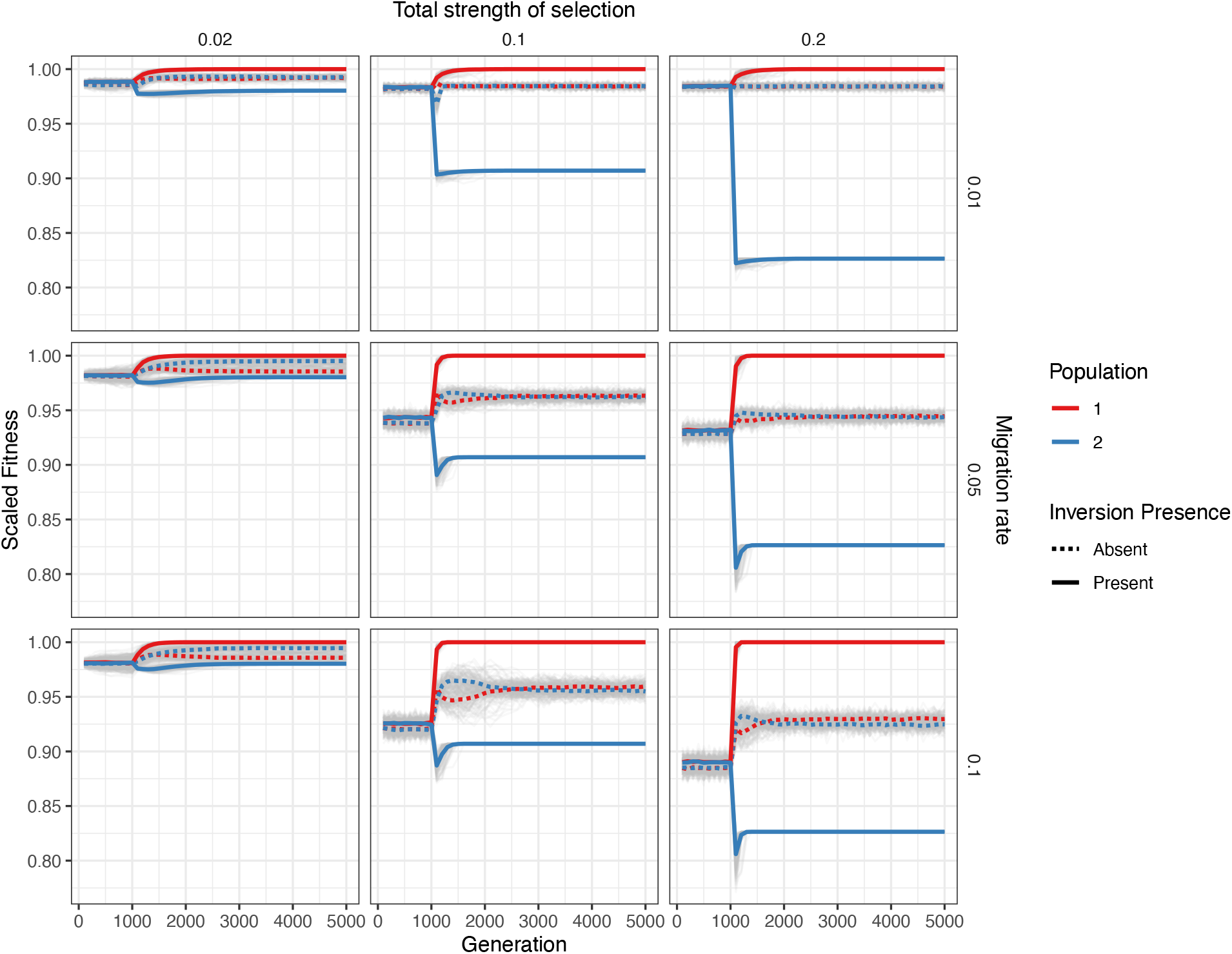
Simulation results for the two populations in the “environmental change” scenario across nine parameter combinations. In this scenario, at generation 1000 the optimal genotype in population 2 shifts to the optimal genotype from population 3 in the “novel environment” scenario described in the text. Each panel depicts the change in scaled mean fitness of a simulated population (colors) over time in the presence or absence of a chromosomal inversion (solid and dashed lines respectively) under a single parameter combination. Each column depicts simulations performed at different total strengths of selection (the sum of the magnitude of all selection coefficients across all loci in any of the given environments). Each row depicts simulations performed at different migration rates. Note that under each parameter combination, the presence of an inversion limits adaptation of population 2 into the changed habitat (i.e., the dashed blue line is always above the solid blue line after generation 1000). It is also interesting to note that population 1 has an increased fitness when the inversion is present, since population 2 contributes less maladaptive gene flow in return to population 1, because it is unable to attain its optimal genotype and has more population 1-adapted genotypes present as compared to without the inversion. For further details on the simulations, see Supplementary Materials.

**Figure S3.**
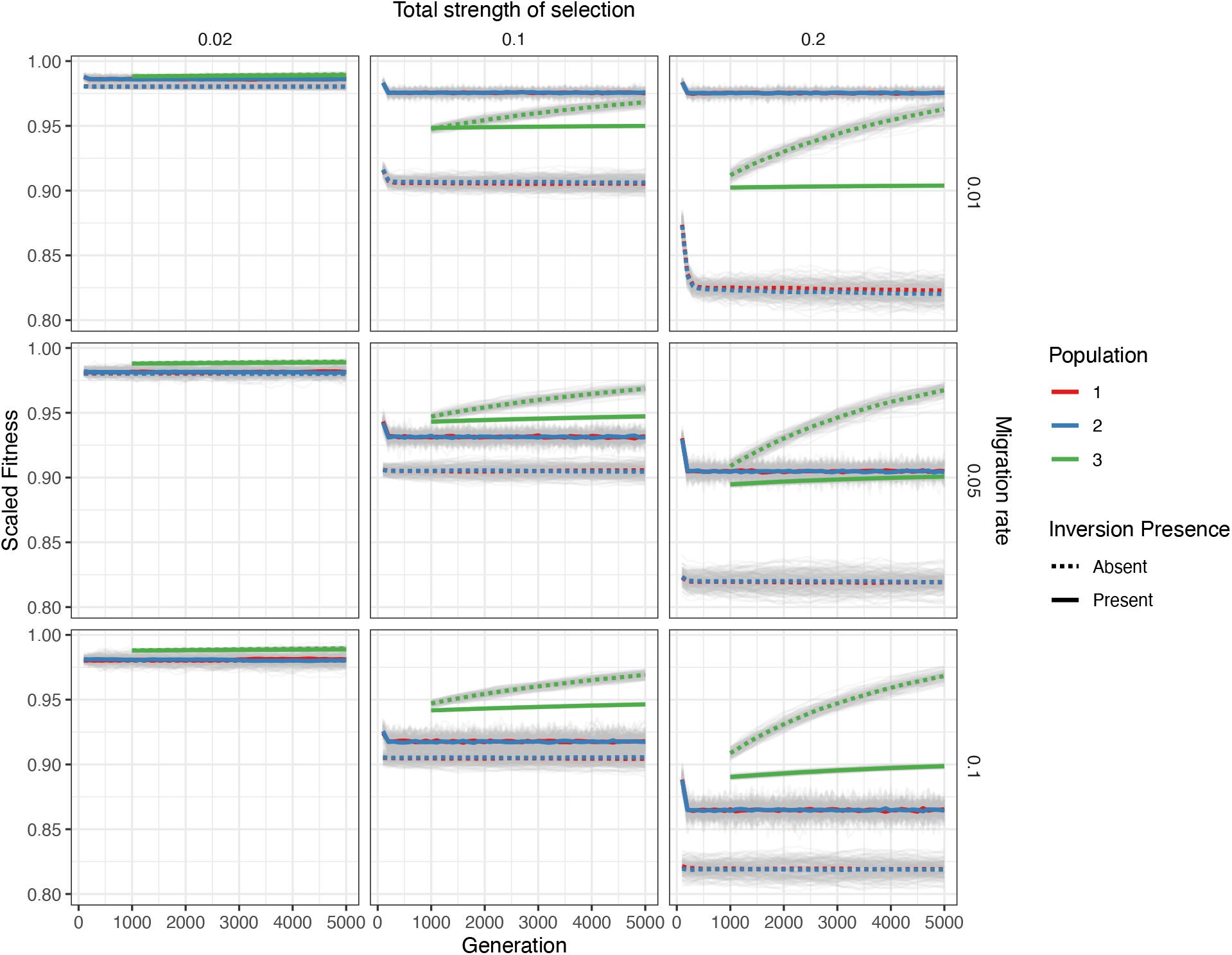
Simulation results for the polygenic scenario (100 loci) across nine parameter combinations. This scenario is identical to the “novel environment” scenario described in the text, except fitness is determined by 100 loci instead of two. Each panel depicts the change in scaled mean fitness of a simulated population (colors) over time in the presence or absence of a chromosomal inversion (solid and dashed lines respectively) under a single parameter combination. Each column depicts simulations performed at different total strengths of selection (the sum of the magnitude of all selection coefficients across all loci in any of the given environments). Each row depicts simulations performed at different migration rates. Note that under all parameter combinations except when the total strength of selection is very weak compared to migration (left panels), the presence of an inversion limits adaptation of population 3 into the novel habitat (i.e., the dashed green line is above the solid green line). We further note that identical simulations but with 40 instead of 100 loci under selection produced highly similar results and are thus not shown.

